# *Escherichia coli* strains from patients with inflammatory bowel diseases have disease-specific genomic adaptations

**DOI:** 10.1101/2021.10.19.464957

**Authors:** Vadim Dubinsky, Leah Reshef, Keren Rabinowitz, Nir Wasserberg, Iris Dotan, Uri Gophna

**Affiliations:** The Shmunis School of Biomedicine and Cancer Research, George S. Wise Faculty of Life Sciences, Tel-Aviv University, Tel Aviv, Israel; The Division of Gastroenterology, Rabin Medical Center, Petah-Tikva, Israel; Felsenstein Medical Research Center, Rabin Medical Center, Petah Tikva, Israel; Sackler Faculty of Medicine, Tel-Aviv University, Tel Aviv, Israel; Colorectal Unit, the Division of Surgery, Rabin Medical Center, Petah-Tikva, Israel

## Abstract

**Objective:** *Escherichia coli* is over-abundant in the gut microbiome of patients with IBD, yet most studies have focused on the adherent-invasive *E. coli* pathotype. Here, we aimed to identify IBD-specific or phenotype-specific genomic functions of diverse *E. coli* lineages.

**Design:** We investigated *E. coli* from patients with UC, CD and a pouch and healthy subjects. The majority of *E. coli* genomes were reconstructed directly from metagenomic samples, including publicly available and newly sequenced fecal metagenomes. Clinical metadata and biomarkers were collected. Functional analysis at the gene and mutation level and genome replication rates of *E. coli strains* were performed, and correlated with IBD phenotypes and biomarkers.

**Results:** Overall, 530 *E. coli* genomes were analysed. A specific *E. coli* lineage (B2) was more prevalent in UC compared to other IBD phenotypes. Genomic metabolic capacities varied across *E. coli* lineages and IBD phenotypes. Specifically, *s*ialidases involved in host mucin utilization, were exclusively present in a single lineage and were depleted in patients with a pouch. In contrast, enzymes that hydrolyze inulin were enriched in patients with a pouch. *E. coli* from patients with UC were twice as likely to encode the genotoxic molecule colibactin than strains from patients with CD or pouch. Strikingly, patients with a pouch showed the highest *E. coli* growth rates, even in the presence of antibiotics. Fecal calprotectin did not correlate with the relative abundance of *E. coli*. Finally, we identified multiple IBD-specific loss-of function mutations in *E. coli* genes encoding for bacterial cell envelope and secretion components.

**Conclusion:** This study presents *E. coli* as a commensal species better adapted to the overly-active mucosal immune milieu in IBD, that may benefit from intestinal inflammation, rather than causing it. The evidence given here suggests adaptive evolution toward attenuated virulence in some *E. coli* strains, coupled with a rapid growth despite the presence of antibiotics.

## INTRODUCTION

Inflammatory bowel diseases (IBD) including Crohn’s disease (CD), ulcerative colitis (UC), and pouchitis are multifactorial diseases characterized by chronic and relapsing intestinal inflammation. CD typically affects the small intestine and colon, UC is confined to the colon, and pouchitis is inflammation of the previously normal small intestine comprising the pouch, in patients with UC after proctocolectomy and ileal-pouch-anal anastomosis [^1^]. The etiology of IBD is believed to involve an aberrant immune response towards the intestinal microbiota [^2^] in a genetically predisposed host [^3^], induced by environmental triggers [^4^].

The gut microbiome in patients with IBD is distinguished by taxonomic [^5,6^] and functional [^7,8^] dysbiosis. One of its hallmarks is the expansion of *Enterobacteriaceae*, in particular of *Escherichia coli* in both fecal [^8^] and mucosal [^9,5^] samples from patients with IBD. *E. coli* is a prevalent commensal in the human gut microbiota [^10^], but has high within species variation in functional potential [^11^], and some strains are pathogens, causing intestinal or extra-intestinal diseases [^12^]. One *E. coli* strain (Nissle 1917) has been considered as a probiotic agent and reported to prolong remission in patients with UC [^13^].

*E. coli* has been extensively studied in the context of IBD pathogenesis, yet the focus has been mostly on the adherent-invasive *E. coli* (AIEC) pathotype [^14,15^]. AIEC strains are characterized by their phenotype of adhesion and invasion into intestinal epithelial cells, and survival and replication within macrophages without prompting cell death [^9^], and are more prevalent in mucosal biopsies of patients with CD compared to patients with UC or healthy controls [^9,16-18^]. Roughly 65% of the AIEC biopsy-derived isolates belong to phylogroup B2 [^16,19^], which contains both commensal and many extra-intestinal pathogenic *E. coli* (ExPEC) strains. Such ExPEC strains typically have pathogenicity islands encoding capsular antigens, iron acquisition systems, cytotoxins and hemolysins [^20^].

Despite considerable efforts, comparative genomic analysis of AIEC strains failed to identify any discriminative genomic markers of this phenotype [^21^]. While most studies investigating *E. coli* as a potential pathogen in IBD focused exclusively on AIEC, the few previous comparative genomic studies of *E. coli* strains in IBD were limited by the number and diversity of the genomes available [^22-24^]. Thus it remains unknown if there are *E. coli* lineages, genes, or alleles that are better adapted to patients with IBD. Here, we performed a large-scale comparative genomic study of *E. coli* from patients with CD, UC and an ileal pouch, aiming to reveal IBD-specific or phenotype-specific adaptations and also test to what degree those strains are expected to promote intestinal inflammation.

## MATERIALS AND METHODS

### E. coli genomes from patients with IBD and healthy subjects and their metadata

The genomes of *E. coli* from patients with a pouch, CD, UC and healthy subjects were obtained from metagenomic and isolate genome data from multiple published studies (supplementary Table S1). In addition, 58 newly sequenced metagenomic samples from 22 patients with a pouch (including two UC samples prior to proctocolectomy and four stoma samples prior to pouch surgery), 9 isolate genomes from patients with a pouch, and 3 isolate genomes from patients with CD from Rabin Medical Center (RMC), Israel, were added to the analysis. Publicly available isolate genomes from patients with CD, UC and healthy subjects and related metadata were downloaded from PATRIC database [^25^]. Publicly available metagenomes used for reconstructing metagenome-assembled genomes (MAGs) were downloaded from the NCBI-SRA repository, and included all phenotypes (supplementary Table S1). In total the comparative genomic analysis included 530 IBD and healthy genomes and 2 *E. coli* Nissle-1917 genomes.

Patients after pouch surgery were recruited at a dedicated pouch clinic in a tertiary IBD referral center in Israel (RMC). The study was approved by the local institutional review board (0298–17) and the National Institutes of Health (NCT01266538). All patients signed informed consent before inclusion. Demographic and clinical data were obtained during clinic visits. Fecal sample collection, fecal calprotectin measurements, genomic DNA extraction and shotgun metagenomic sequencing, pouch disease behavior (phenotype) definition and treatment protocol were performed as described in [^26^].

### Pipeline for the reconstruction of genomes from metagenomes

To reconstruct the genomes of *E. coli* directly from metagenome samples (*E. coli* MAGs), we applied the following single-sample metagenomic assembly pipeline to each sample. (i) Metagenome reads were quality-filtered with Trimmomatic v0.36 [^27^] using default parameters, removing low-quality and short reads, low-quality bases, and clipping Illumina adaptors. Human-derived reads were removed by mapping them against the human genome (GRCh38) with Bowtie2 v2.2.9 [^28^] in --very-sensitive-local mode. (ii) The remaining reads were mapped against a reference database of the *E. coli* pangenome to retain only closely matching reads using MIDAS [^29^]. (iii) Samples with a sufficient amount of mapped reads to *E. coli* pangenome (depth of coverage >5x) were separately *de-novo* assembled with Unicycler v0.4.4 [^30^], using default parameters. Contigs shorter than 1 Kb were discarded. (iv) To obtain better quality MAGs, the remaining contigs were screened with BLASTN against the NR database, and only contigs that aligned to *E. coli* genes as top hit with nucleotide identity of ≥90% were retained. (v) All the MAGs underwent the following quality control steps: genome completeness and contamination measures were assessed using CheckM v1.0.7 [^31^] in taxonomy-specific mode (taxonomy_wf) for the species *E. coli*. MAGs with levels of completeness <75% and contamination >5% were discarded. The amount of strain-level heterogeneity was estimated with CMSeq (https://github.com/SegataLab/cmseq), which calculates the polymorphism at each position (with coverage of >5x) in the contigs (a position was considered as non-polymorphic if the dominant allele frequency was >80%). (vi) The relative abundance of each MAG from the corresponding metagenome sample was calculated by mapping reads against the reconstructed MAG from the same sample using Bowtie2 (--very-sensitive mode), and to avoid counting reads from closely related strains, only aligned reads with an edit-distance (mismatches) of **≤** 2 bases to the contig were considered, divided by the total number of quality-filtered reads in the sample.

In total 412 *E. coli* MAGs were reconstructed and had genome completeness of >75% and contamination <5%, while 346 MAGs could be defined as high-quality according to current guidelines with completeness of >90% [^32^] - see supplementary Table S2 for assembly quality control statistics.

### Whole-genome phylogeny and average nucleotide identity analysis

Phylogenetic analysis of the *E. coli* genomes (MAGs and isolates) was performed using PhyloPhlAn-3 [^33^]. The phylogeny was built by annotating the input genomes with a set of species-specific marker genes identified from the UniRef90 database (2818 *E. coli* markers). The following parameters were used, ‘--diversity low --accurate -min_num_entries 505 -- force_nucleotides’, to generate high-resolution strain-level phylogeny (nucleotides level) with the genes defined as core if present in ≥95% of the genomes.

Whole-genome average nucleotide identity (ANI) between pairs of genomes was calculated with FastANI v1.32 [^34^] using default parameters. ANI values of ≥95% are the most frequently used as a species-level taxonomic boundary threshold for prokaryotic genomes [^34^].

### Functional annotation and analysis of the genomes

Functional annotation of the *E. coli* genomes was performed based on several tools and databases. EggNOG mapper v2.0.1 [^35^] was used to generate functional gene profile (pangenome) based on EggNOG orthology system with DIAMOND v0.9.24 [^36^] for the sequence similarity search. In addition, the genomes were annotated with KEGG Orthologs (KO) assignment based on hidden Markov model profiles using KofamScan v1.3.0 [^37^]. Carbohydrate-active enzymes (CAZy) families [^38^] were identified using ‘hmmscan’ from HMMER v3.2.1 (http://hmmer.org) against dbCAN HMMs database release-9, applying stringent filtering cutoff as suggested by the authors (http://bcb.unl.edu/dbCAN2). To add virulence genes annotation to the genomes, protein search Blastp (DIAMOND) against the core set of experimentally verified virulence factors database (VFDB) was performed [^39^]. Only sequences that had identity ≥ 90% to the proteins annotated in VFDB were considered. Similarly, antibiotic resistance genes were identified by using the curated protein sequences (homolog model) in the comprehensive antibiotic resistance database (CARD) [^40^]. Only sequences that had identity ≥ 95% to the proteins in CARD were considered. For all the functional annotations, protein-coding genes within the contigs were predicted with Prodigal v2.6.3 and used as input query for sequence similarity searches.

### Genome replication rates of E. coli within subjects

Genome replication rate for each *E. coli* MAGs was calculated with the iRep (genome replication index) algorithm [^41^]. This tool is based on the principle that actively replicating cells have higher coverage near the origin of replication, compared to the terminus in their genomes [^42^]. Only MAGs which were obtained from fecal and aspirate metagenomes were used in this analysis (n=391), since the mapping of metagenome reads to the assembled MAGs is required for this tool.

### Mutational analysis of E. coli functional genes

To identify IBD-specific adaptations at the mutational level, Snippy v4.6.0 variant calling tool (https://github.com/tseemann/snippy) was used. The analysis was performed in a within-clade approach with a different isolate reference genome (high-quality draft) for each *E. coli* clade as was determined by the whole-genome phylogeny. For clade-I, clade-III, clade-IV and clade-V, the following genomes from healthy subjects were used as reference (PATRIC genome ID), 749549.3, 749550.3, 749540.3, 749545.3, respectively. Functional annotations for these genomes were obtained from NCBI-Assembly according to NCBI prokaryotic genome annotation pipeline in genomic GenBank format, as required by Snippy. The analysis was not performed for clade-II due to the small number of genomes. The metagenomic reads composing the MAGs from IBD patients were mapped against the reference genomes, with a minimum read depth at a position for variant calling was set to 10 (default). Mutation variants were identified in coding and non-coding regions in the genome, and were classified as synonymous, non-synonymous, insertions/deletions and frameshifts. Synonymous mutations and mutations in intergenic regions were not analyzed further.

### Statistical analysis

All statistical analysis was performed in R v3.6.3. The Kruskal-Wallis H-test with Dunn’s test correction for multiple pairwise comparisons was performed using ‘kruskal.test’ and ‘dunn.test’ functions. Wilcoxon (Mann-Whitney) rank test was performed using the ‘wilcox.test’ function. Fisher’s exact test with false discovery rate correction (Benjamini-Hochberg) for multiple hypothesis testing was performed with the ‘fisher.test’ and ‘p.adjust’ functions. Spearman’s rank correlation was performed with the ‘cor.test’ function. Permutational multivariate analysis of variance (PERMANOVA) test was performed using the *adonis* function. A linear mixed-effects model of iRep (dependent variable) as a function of genome and patient related predictor variables, with individual patients from which a set of samples were derived being specified as a random effect, was performed with the ‘lmer’ function.

### Data availability (Data sharing statement)

All the metagenomic sequence data generated in this study (patients with a pouch and UC and a stoma) have been deposited in NCBI-SRA and the isolate genomes of patients with a pouch and CD were deposited in NCBI-Genome, and are available under BioProject number PRJNA730677. The assembled MAGs are available for download from Figshare data repository with the following URL: https://doi.org/10.6084/m9.figshare.14625537.

## RESULTS

We studied the genomes of over 530 *E. coli* strains from fecal (n=438), aspirate (n=24) and biopsy (n=68) samples of patients with IBD (n=459) and healthy individuals (n=71). We established a dataset that encompassed 21 individual studies, both publicly available and newly sequenced samples from our own cohort of patients with a pouch (supplementary Table S1). We reconstructed the majority of the analyzed genomes (77%) directly from metagenomic samples (metagenome-assembled genomes [MAGs]), while the rest of the genomes were obtained from cultured isolates. The patients with IBD in this study from which we obtained *E. coli* genomes included the following clinical phenotypes: Pouch (n=268), CD (n=124), UC (n=63) and additional four patients with UC and a stoma (prior to pouch surgery). For a subset of patients from our dataset, we had fecal calprotectin and antibiotic usage data (supplementary Table S1).

### *E. coli* strains from patients with IBD and healthy subjects belong to distinct lineages that are differentially distributed across phenotypes

First, the phylogenetic structure of *E. coli* strains was investigated using whole-genome analysis of core marker genes. We identified five distinct clades comprising the *E. coli* strains in our dataset (Fig. 1A), which we named Clade-I to Clade-V. Each clade was supported by isolate genomes intermixed with the MAGs, reassuring that our computational approach for MAGs reconstruction from metagenomes is robust, and the MAGs are compatible with isolate genomes. Although the majority of the *E. coli* in our dataset were from fecal samples, the genomes obtained from biopsies and aspirates clustered with the fecal strains throughout the five clades, implying that there is no clade that is specifically mucosa-associated.

**Figure 1.**
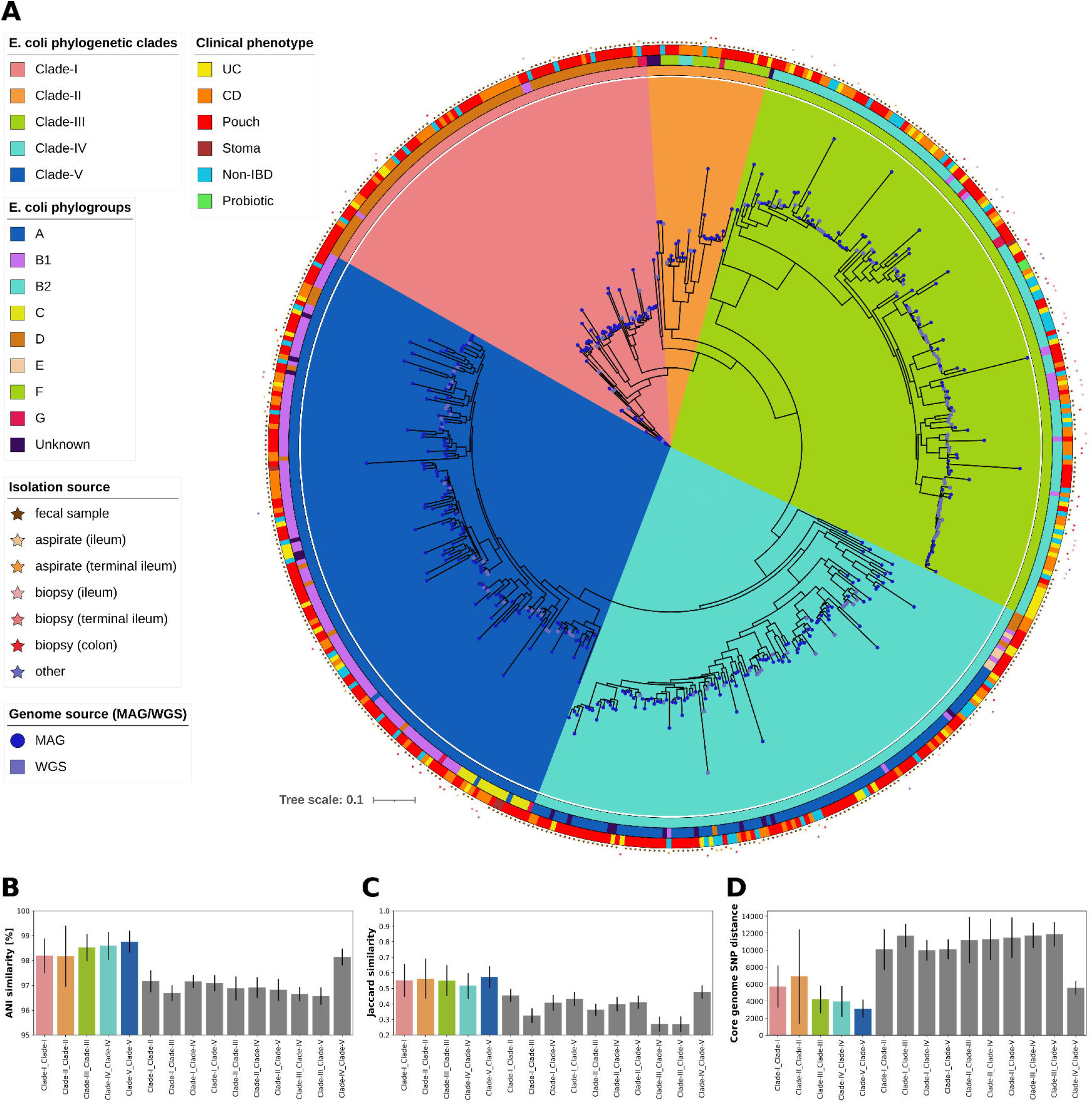
Phylogenetic structure of the metagenome-assembled genomes (MAGs) and isolate genomes of *E. coli* from patients with IBD and healthy subjects. (**A**) A whole-genome phylogenetic tree based on core genes of 532 *E. coli* genomes. The inner ring indicates the clinical phenotype of the subject from whom the genome was derived; the middle ring corresponds to the *E. coli* phylogroup membership; the outer ring is the *E. coli* clade membership based on whole-genome phylogeny; circles/squares on tree nodes denote whether the genome is a MAG or isolate genome (WGS label), respectively; stars outside the rings denote the isolation source of the sample. The phylogenetic tree was built with RAxML (parameters “-p 1989 -m GTRCAT”) as part of PhyloPhlAn-3 tool [Asnicar et al. 2020], based on 2818 species-specific marker genes defined as core, that were present in ≥95% of the genomes. The branch scale is nucleotide substitutions per site. (**B**) Whole-genome average nucleotide identity (ANI) between pairs of *E. coli* genomes within a clade and between the clades. (**C**) Jaccard similarity in gene content (non-core functional genes based on EggNOG) between pairs of *E. coli* genomes within a clade and between the clades. (**D**) Core genome single-nucleotide polymorphism (SNP) distance between pairs of *E. coli* genomes within a clade and between the clades. In panels **B** to **D**, colored bars are within a clade and gray bars are between the clades comparison, respectively.

Population genetic studies of *E. coli* isolates have traditionally relied on the division into eight deep branching lineages called phylogroups [^43^]. An analysis of *E. coli* phylogroups (Fig. 1A) indicated that the IBD strains originated from a diverse array of lineages (A, B1, B2, C, D, E, F, G). The five clades we identified based on whole-genome largely coincided with the phylogroups, with clade-I being consistent with D, clade-II with F, clade-III with B2, clade-IV with A and clade-V with B1 and C. Clade-III to clade-V were represented by the largest and comparable number of genomes (n=146, 126, 143), clade-I was smaller (n=84), and clade-II was represented by only 27 genomes. The separation of *E. coli* into five distinct clades was further supported based on higher average nucleotide identity (ANI) within-clade (mean of 98.17% to 98.75%) compared to a mean ANI similarity of 96.56% to 98.14% between clades (Fig. 1B), as well as by gene content and core genome SNP distance (Fig. 1C-D).

We next analyzed the distribution of the five *E. coli* clades among the clinical phenotypes. All the IBD phenotypes and healthy subjects had at least one strain from each clade, suggesting that many different *E. coli* lineages successfully colonize both patients with IBD and healthy subjects. Nonetheless, we observed an uneven distribution in clade-III (Fisher’s exact test, FDR p < 0.05; Fig. 2A), which was more prevalent in UC (49.2%) and non-IBD (40.8%) compared to CD (31.5%) and pouch (17.5%), whereas clade-I prevalence in UC was only 4.8%, the lowest among the phenotypes.

**Figure 2.**
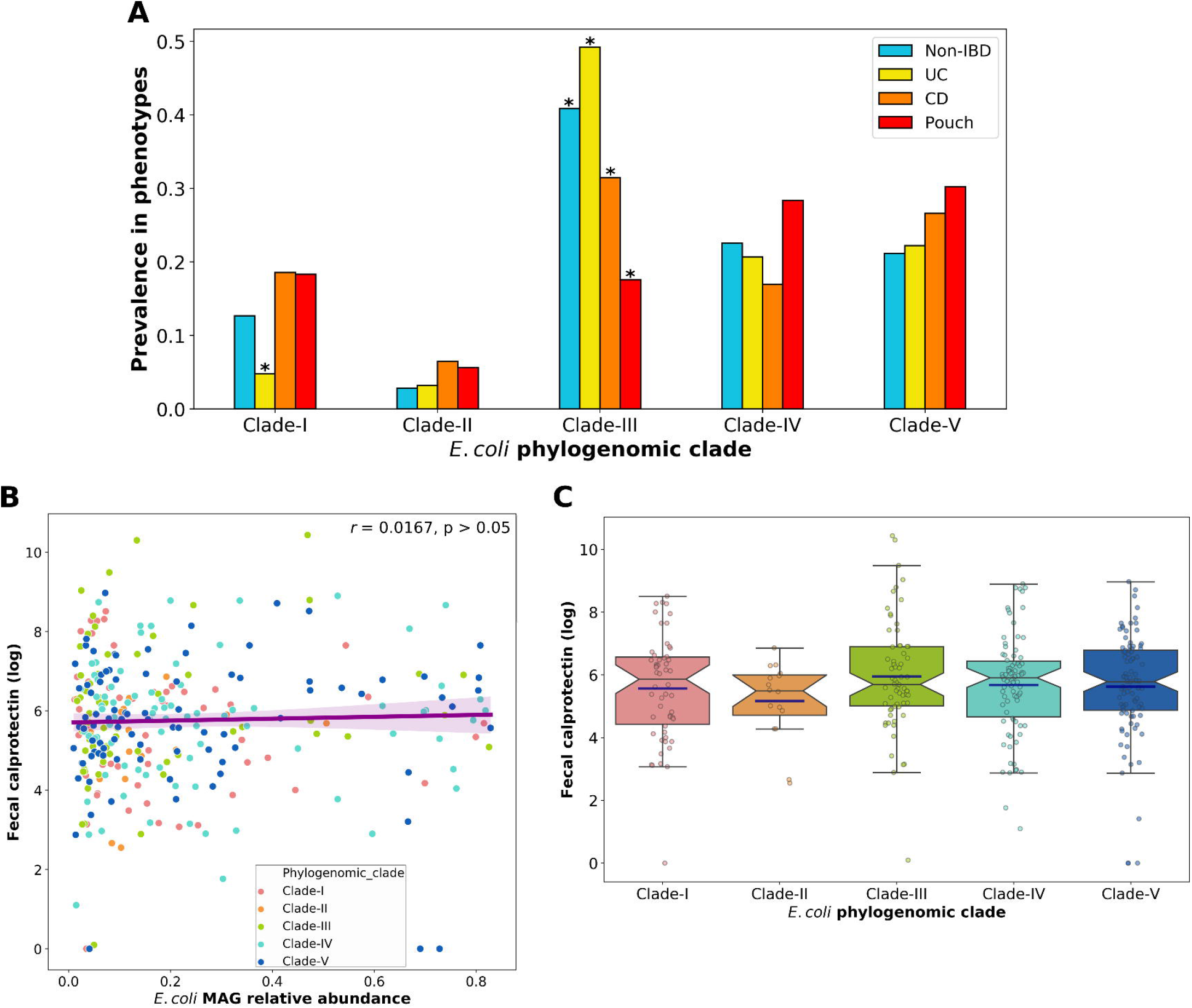
Distribution of the *E. coli* clades across clinical phenotypes and lack of association between intestinal inflammation and *E. coli* clade and relative abundance. (**A**) Prevalence of *E. coli* clades in the different clinical phenotypes of patients with IBD (pouch, CD and UC) and non-IBD subjects. *FDR p < 0.05; Fisher’s exact test. (**B**) Spearman correlation between *E. coli* MAGs relative abundance and fecal calprotectin level (log-transformed). (**C**) Fecal calprotectin levels (log-transformed) associated with each *E. coli* clade. Box plot whiskers mark observations within the 1.5 interquartile range of the upper and lower quartiles. The blue horizontal lines mark the mean value in each clade.

Finally, we tested to what degree can *E. coli*, or specific clades of this species be associated with intestinal inflammation. Fecal calprotectin, used as a biomarker of intestinal inflammation, did not correlate with the relative abundance of the *E. coli* MAGs in the microbiomes from corresponding metagenomic samples. (Spearman r = 0.0167, p > 0.05; Fig. 2B). Furthermore, the median calprotectin levels associated with each clade were similar (Kruskal-Wallis, p > 0.05; Fig. 2C), although in the group of patients with clade-III *E. coli* the mean calprotectin was higher due to several patients with very high intestinal inflammation (> 8000 mcg g^-1^).

#### E. coli clades present in patients with IBD may have particular functional potential

We next explored the functional potential of the *E. coli* clades through genome annotation based on several databases (EggNOG, KEGG orthologs [KO], carbohydrate-active enzymes [CAZy], virulence factors and antibiotic resistance databases). We identified a pangenome of 7,462 genes according to the EggNOG (with either a known gene name or KO number), of which 2,121 were non-core (present in less than 90% of the genomes) and thus might drive differences in the functional potential between the clades. A substantial variation in the functional genes (EggNOG, non-core) was observed between the *E. coli* genomes (Fig. 3A), which was explained to a large degree by the clades membership (PERMANOVA test, R^2^ = 0.437, p < 0.001). Clade-III was the most dissimilar from the other clades, while clades IV and V had some degree of overlap. A similar clustering pattern was observed according to KO and CAZy gene annotations (Fig. 3B, C) and also based on core-gene SNP alignment (Fig. 3D). Focusing on genes with KO annotations, overall we found 709 non-core genes that significantly differed in prevalence between the *E. coli* clades (Fisher’s exact test, FDR p < 0.05; supplementary Table S3). Many genes shared a similar prevalence pattern between clades IV and V and were nearly absent from clade-III, and on the other hand, clade-III was enriched in genes that were absent from other clades.

**Figure 3.**
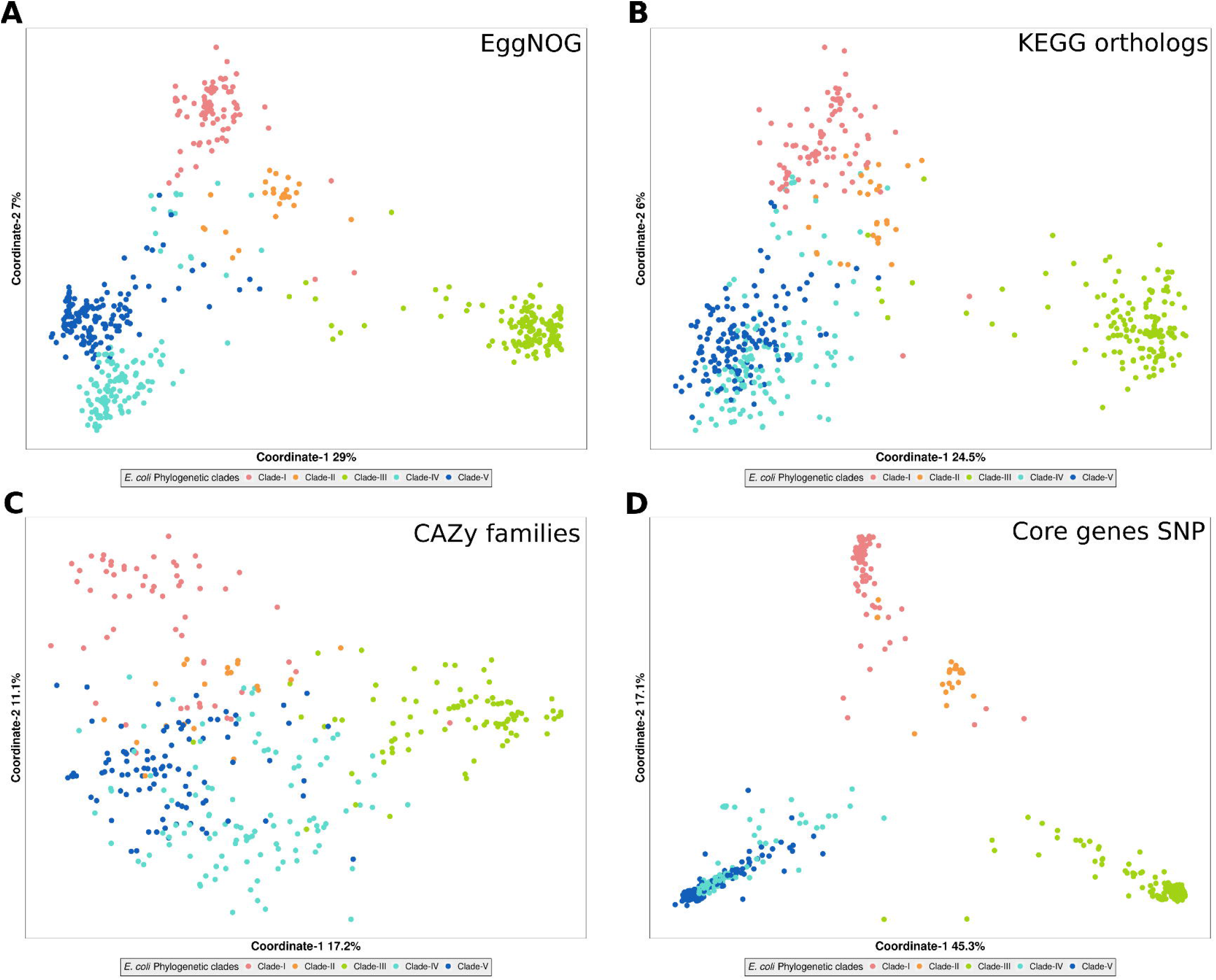
Variation in the functional gene content of the *E. coli* genomes explained by clade membership. (**A**) Principal coordinates analysis (PCoA) of Jaccard distances (overall dissimilarity in the presence or absence of genes) based on non-core genes annotated with EggNOG. (**B**) PCoA of Jaccard distances based on non-core genes annotated with KEGG Orthologs (KO). (**C**) PCoA of Jaccard distances based on non-core carbohydrate-active enzymes (CAZy) families. (**D**) PCoA based on dissimilarity in core-gene single-nucleotide polymorphisms (SNPs). The variation in the functional gene content explained by the clade membership of *E. coli* genomes in panels **A** to **D** (PERMANOVA, R^2^ = 0.437, 0.401, 0.366 and 0.748, respectively; p < 0.001).

We also specifically analyzed the CAZy families to identify potential differences in carbohydrate utilization between the clades. Even though many of the identified 26 non-core CAZy families were broadly distributed, we nonetheless observed a distinct prevalence pattern between the clades for some of them (supplementary Table S4). The GH38 CAZy of alpha-mannosidase (involved in the oligosaccharide mannose degradation) was over 93% prevalent in clades IV and V, but less than 33% prevalent in the other three clades (Fisher’s exact test, FDR p < 9.6 × 10^−63^). On the other hand, the GH127 CAZy family involved in other oligosaccharides degradation including arabinofuranose was 98.8% and 100% prevalent in clades I and II, while in the other clades it was only 29.5% to 67% prevalent. Unlike the other clades, clade-III almost exclusively possessed a high prevalence (98%) of GH33 sialidases (Fisher’s exact test, FDR p < 3.4 × 10^−113^), which are involved in the breakdown and utilization of mucin-derived human glycans (by cleaving terminal sialic acid from sialylated mucins; ^44^).

#### E. coli clades present in patients with IBD have distinct virulence factors and antibiotic resistance genes repertoires

To determine whether some *E. coli* clades in our dataset have a higher virulence potential and carry more antibiotic resistance genes than others, we screened the genomes based on VFDB (supplementary Table S5) and CARD (supplementary Table S6) databases, respectively (included only non-core genes, as defined above). We noted a clear difference in the virulence factors profile between the clades, with clade-III (B2 phylogroup equivalent) genomes encoding a higher number of potentially proinflammatory genes and toxins (Fig. 4A). Half of the genomes in clade-III had genes encoding for colibactin (pks genomic island - *clb*), a genotoxic molecule that can induce double-strand DNA breaks, eukaryotic cell cycle arrest, and chromosome aberrations [^45^]. Several serine-protease autotransporters that are harmful for eukaryotic cells [^46^] and are considered as enterotoxins (*pic, sat, vat*), were almost exclusively present in clade-III genomes, and their prevalence ranged from 30% to 77%. Moreover, *α*-hemolysin (*hly*) which was shown to disrupt epithelial cell tight junctions in patients with UC and thus promote intestinal inflammation [^47^] was present in 25% of clade-III genomes. Notably, almost none of the genomes from the other clades possessed the above mentioned virulence factors (Fig. 4A).

**Figure 4.**
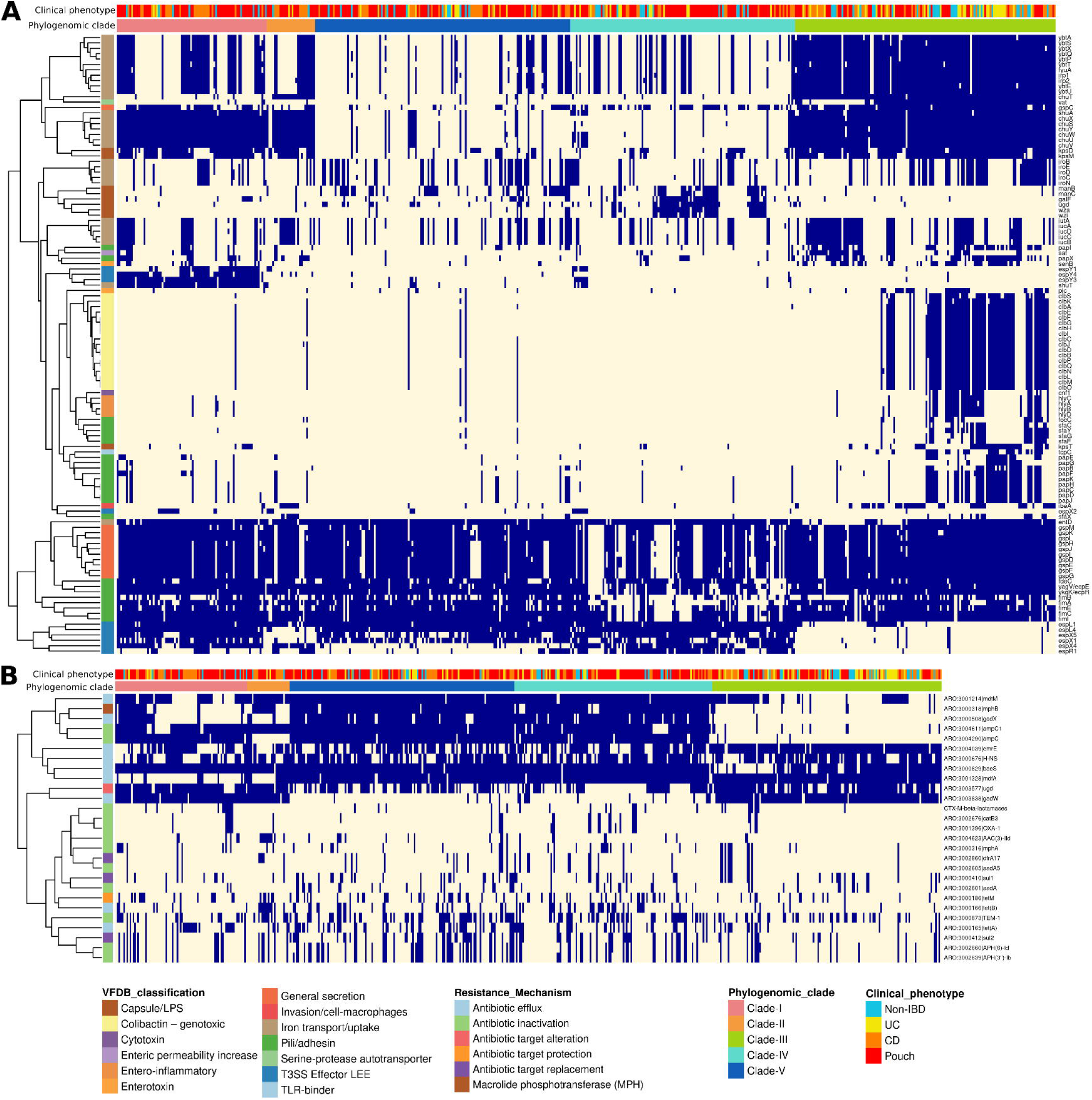
Virulence factor and antibiotic resistance gene profiles in the *E. coli* genomes from patients with IBD and healthy subjects. (**A**) Heatmap of virulence factors (rows) in *E. coli* genomes (columns) showing the presence (blue) or absence (beige) of the genes. (**B**) Heatmap of antibiotic resistance genes (rows) in *E. coli* genomes (columns) showing the presence (blue) or absence (beige) of the specific genes. Color bars at the top of the plots represent the clinical phenotype of the subject and the clade membership of the *E. coli* genomes. Color bars at the left of the plots represent the classification category of the virulence genes in panel (**A**) and the antibiotic resistance mechanism in panel (**B**). Genes were clustered using euclidean distances by applying the complete linkage method. For the list of p-values obtained from Fisher’s exact test for genes prevalence, see supplementary Table S5 and Table S6.

Genes for synthesis of capsular polysaccharides were enriched in clades IV and V, whereas clades I to III shared a high prevalence (92% to 100%) of genes related to iron transport (heme-binding genes cluster *chu* and *shu*). Interestingly, clade-III genomes contained an additional iron uptake system (Fisher’s exact test, FDR p < 1.5 × 10^−44^), namely yersiniabactin encoded on a pathogenicity island, which is also involved in copper uptake [^48^]. Lastly, we observed that 16 genes coding for pili that enable extra-intestinal colonization (*pap* and *sfa*), were more abundant in clade-III, but had a relatively low mean prevalence of 28% (Fig. 4A).

We also explored the antibiotic resistance genes in the *E. coli* strains from each clade. Genes encoding various efflux pumps (against tetracycline, macrolide, aminocoumarin and aminoglycoside antibiotics) showed a diverse prevalence pattern between the clades (Fig. 4B). Aminoglycoside resistance by inactivation had a relatively low prevalence in all the clades (< 37%), but *AAC(3)-IId* was significantly higher in clade-II than in the other clades (Fisher’s exact test, FDR p < 2.2 × 10^−6^). We also detected beta-lactamase encoding genes, some of which were prevalent (*ampC* and TEM-1) while others were relatively rare, especially those coding for the CTX-M family of extended-spectrum *β*-lactamases [^49^]. Interestingly, genomes from clade-III had on average the lowest number of antibiotic resistance genes (Fig. 4B).

For 408 *E. coli* genomes, we obtained information regarding the antibiotic usage of the patients that the samples were obtained from. By dividing these genomes into two groups, from subjects that received antibiotic treatment in the last month (Abx+) and those that were free of antibiotics for at least one month (Abx-), we could detect 14 antibiotic resistance genes that were significantly enriched in the genomes from Abx+ group (Fig. 5A). Importantly, *aac(6’)-Ib-cr, oxa-1* and *catB3* genes conferring resistance against fluoroquinolones, beta-lactams and chloramphenicol, respectively, were 16% prevalent in the genomes of Abx+, compared to only 2.54% in the Abx-(Fisher’s exact test, FDR p < 8.4 × 10^−5^). These genes are known to be encoded together on plasmids of *Enterobacteriaceae* [^50^], and accordingly, we observed these genes co-located on the same contigs of *E. coli* in our dataset. Overall, genomes in the Abx+ and Abx-groups had a median of 11 and 9 resistance genes, respectively (Mann-Whitney, p < 5.3 × 10^−4^; Fig. 5B), suggesting that antibiotic treatment increases mobile antibiotic resistance in *E. coli*.

**Figure 5.**
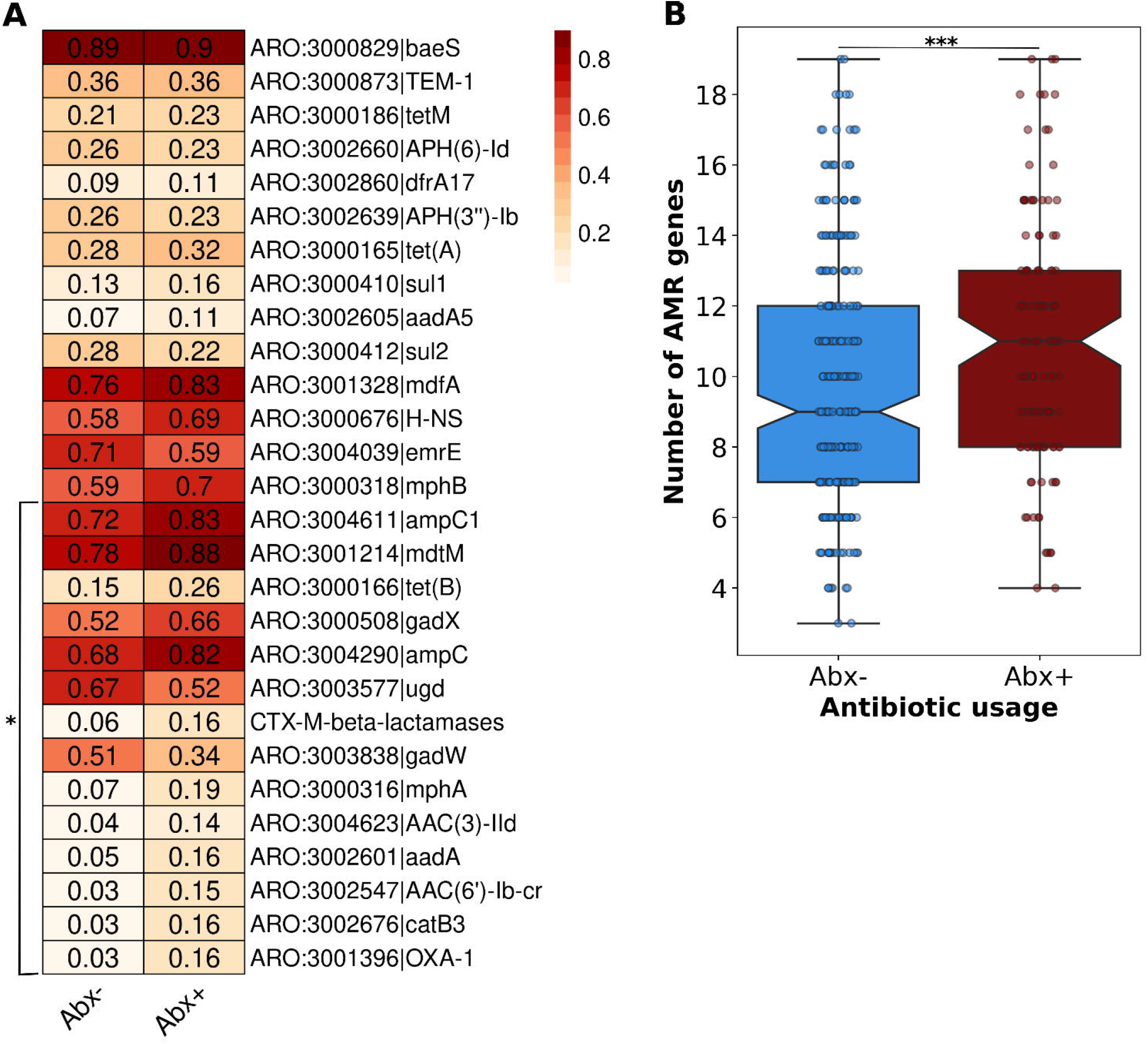
Higher prevalence of antibiotic resistance genes of *E. coli* in subjects undergoing antibiotic treatment. (**A**) Heatmap showing the prevalence of antibiotic resistance genes in *E. coli* genomes from subjects that received antibiotic treatment in the last month (Abx+) compared to those that were free of antibiotics for at least one month (Abx-). *FDR p < 0.05; Fisher’s exact test. (**B**) The distribution of the number of antibiotic resistance genes (AMR) in *E. coli* genomes from Abx+ and Abx-subjects as defined in panel **A**. *p < 0.001; Mann-Whitney test. Box plot whiskers mark observations within the 1.5 interquartile range of the upper and lower quartiles.

#### Genome replication rates of E. coli strains are associated with clinical phenotypes but not with intestinal inflammation or antibiotic usage

To test how well do *E. coli* strains grow in the IBD gut, we analyzed the growth dynamics of the *E. coli* genomes obtained from 375 fecal and 16 aspirate metagenomes of patients with IBD and healthy subjects *in vivo* using iRep (genome replication index [^41^]). Patients with a pouch had the fastest-growing *E. coli* strains (Kruskal-Wallis, p < 0.05; Fig. 6A), followed by patients with CD, while those with UC and non-IBD subjects had the slowest-growing strains. It is worth noting that iRep was not correlated with the relative abundance of *E. coli* (Spearman r = -0.018, p > 0.05; Fig. 6B). A linear mixed-effects model of iRep taking into account strain and patient-related parameters indicated that only the clinical phenotype of the subject was significantly associated with iRep (mixed-effects model, p < 0.001). In contrast, the phylogenetic clade and relative abundance of *E. coli*, antibiotic usage (antibiotics in the last month) and fecal calprotectin (intestinal inflammation marker) failed to show statistically significant associations with iRep.

**Figure 6.**
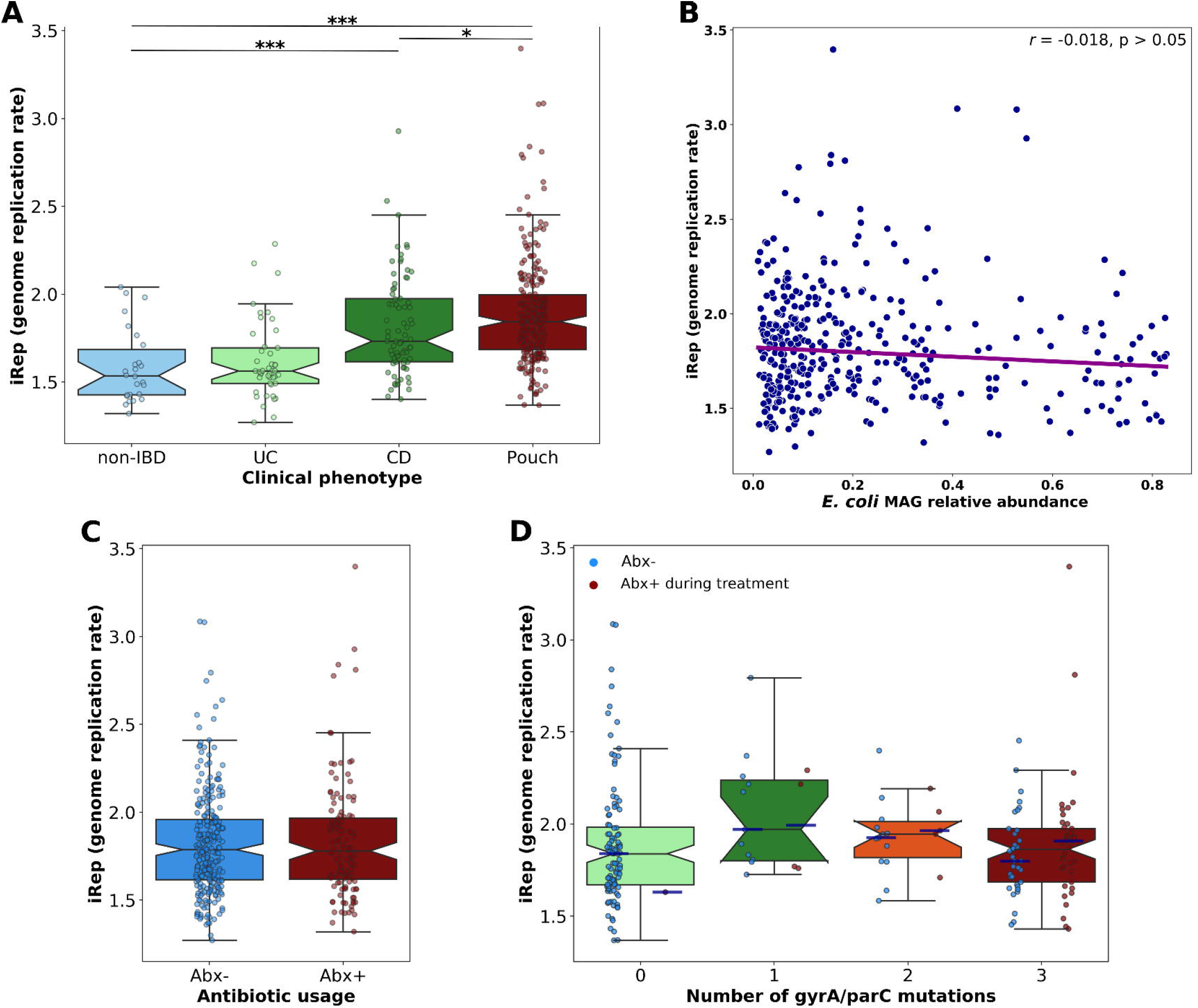
Inferred growth rates of *E. coli* strains from patients with IBD and healthy subjects. (**A**) Inferred growth rates (using iRep [genome replication index]) of *E. coli* strains from patients with a pouch, CD, UC and healthy subjects. *p < 0.05; ***p < 0.001; Kruskal-Wallis test. (**B**) Spearman correlation between iRep and the relative abundance of *E. coli* in the microbiomes from corresponding metagenomic samples. (**C**) iRep of *E. coli* strains from subjects that used antibiotics in the last month (Abx+) compared to subjects who were antibiotic-free for at least one month (Abx-). (**D**) iRep of *E. coli* strains from patients with a pouch that were treated with ciprofloxacin, grouped according to the number of resistant mutations in drug target genes (*gyrA* and *parC*). Zero mutations represent ciprofloxacin sensitive strains. “Abx+ during treatment” and “Abx-” division define samples from patients taken during antibiotic treatment or in the absence of antibiotics for at least one month, respectively. In panels **A, C**, and **D**, box plot whiskers mark observations within the 1.5 interquartile range of the upper and lower quartiles.

Curiously, *E. coli* strains that grew in the presence of antibiotics had similar iRep to those from non-antibiotic treated patients (Mann-Whitney, p > 0.05; Fig. 6C). For a subset of patients with pouch (N=233), for which we had more comprehensive antibiotic usage data that were treated with ciprofloxacin, we could match resistance mutations with the respective iRep from the same sample. As the main mechanism of resistance again ciprofloxacin (fluoroquinolone) is point mutations in drug target genes, *gyrA* and *parC* [^26^], we extracted these genes and analyzed them for mutations in previously known positions conferring resistance. Even taking into account *E. coli* from samples taken during active antibiotic treatment, highly resistant strains with 3 mutations had a similar iRep to sensitive strains (no mutations) [Fig. 6D], suggesting that the resistant strains grow very well within hosts, with rates comparable to sensitive strains growing in the absence of antibiotics.

#### IBD-associated functional genes within the Clade-III (B2 phylogroup) lineage

Our next goal was to determine if there is an association between specific functional genes of *E. coli* and patients with IBD (Pouch, CD and UC phenotypes) or healthy subjects. As we have shown above, many functions (including metabolism-, virulence- and resistance-related) exhibit strong clade-dependent distribution patterns. Thus, in order to overcome such biases, we looked at within-clade functional differences between various IBD phenotypes and healthy controls. Focusing on clade-III, we observed 154 non-core KO annotated genes that differed significantly in prevalence between the phenotypes (Fisher’s exact test, FDR p < 0.05; supplementary Table S7). Specifically, the gene *scrK* encoding fructokinase (fructose metabolism) was highly enriched in patients with a pouch (Fisher’s exact test, FDR p < 1.25 × 10^−6^). Moreover, sucrose and the fructose-containing polysaccharide inulin degrading CAZy family (GH32) was enriched in patients with a pouch, with a prevalence of 85%, compared with 35% to 54% in the other phenotypes (Fisher’s exact test, FDR p < 1.15 × 10^−3^). In contrast, patients with a pouch were depleted (6.4%) in endo-alpha-sialidase (GH58; involved in utilization of sialylated mucins), which was significantly more abundant in healthy subjects (27.6%), and patients with CD (25.6%) and UC (45%) [supplementary Table S8]. In addition, genes involved in manganese and iron transport (*sitA*-*D*) were more abundant in UC and healthy than in pouch and CD phenotypes (Fisher’s exact test, FDR p < 0.05; supplementary Table S7). Finally, we did not find significant differences in the prevalence of propanediol dehydratase (*pduC*) between patients with CD and healthy subjects, in contrast to a recent report [^51^].

In terms of virulence factors, the colibactin gene cluster was significantly more abundant in patients with UC (66.5%) and healthy subjects (58.6%) than in patients with CD (28.2%) and a pouch (36%). The serine-protease *pic* had a prevalence of 47% in *E. coli* from patients with a pouch but was less than 20% prevalent in the other phenotypes. Surprisingly, healthy subjects were significantly enriched in *senB* and *cnf1*, encoding for enterotoxic and cytotoxic proteins, respectively. Finally, *pap* genes, which enable adhesion to urinary tract epithelial cells, were twice as abundant in *E. coli* from healthy subjects and patients with UC compared to CD and patients with pouch (supplementary Table S9). The only antibiotic resistance gene that was differently prevalent was *tetM*, conferring resistance against tetracyclines, which was solely found in 30% of the genomes in patients with a pouch (Fisher’s exact test, FDR p < 3.3 × 10^−5^; supplementary Table S10).

#### Mutational analysis of functional genes reveal IBD-specific adaptations

To complement the analysis of the differential prevalence of functional genes in patients with IBD and healthy subjects, we looked for whether there are IBD-specific adaptations at the mutational level. We mapped the metagenome reads composing the MAGs from IBD phenotypes against a reference isolate genome from the same clade from a healthy phenotype, richly annotated by the NCBI prokaryotic genome annotation pipeline. We defined IBD-associated mutations as those appearing in over 30% of IBD-derived genomes and absent from all non-IBD strains within a clade. We identified a large number of non-synonymous mutations in functionally-annotated genes of *E. coli* strains from clade-III (supplementary Table S11), of which 126 mutations were present in over 50% of the analyzed genomes, and 639 occurred in at least 30%.

Remarkably, some genes were especially mutation-prone, showing independent multiple mutations in the same gene or operon in different genomes, or function-inactivating mutations such as frameshifts (Table 1). Such convergent mutational patterns were especially notable in genes encoding for bacterial cell surface- or secretion-associated components. These included flagellar assembly and secretion proteins (*flhABE*, including a frameshift mutation in *flhB, and fliFH*), colanic acid biosynthesis (*wcaCLM*), outer membrane proteins OmpA and AsmA and the intimin-like adhesin FdeC. Such proteins may have direct proinflammatory roles in the gut, and some of them are known to be important antigens [^52-54^]. Interestingly, the gene encoding the lipoprotein metalloprotease SslE, which contributes to activation of macrophages via toll-like receptor 2 [^55^], showed seven different mutations, including two frameshifts. Moreover, the type II secretion system which is responsible for the secretion of SslE, had 11 mutations in its encoding operon (Table 1). In addition, yersiniabactin non-ribosomal peptide synthetase (*irp*2) which is part of the yersiniabactin iron uptake system also showed four different mutations. Other frequently mutated genes with a possible role in virulence included the type VI secretion system [^56^], with mutations in several genes from the cluster (*tssHGK* and *vgrG*). To confirm that those mutations could alter the antigenicity of the respective proteins, we mapped them onto the three-dimensional structure of the respective proteins, using ConSurf [^57^]. This analysis revealed that 81% of the mutations in those genes indeed were in the surface-exposed residues (Table 1).

**Table 1.**
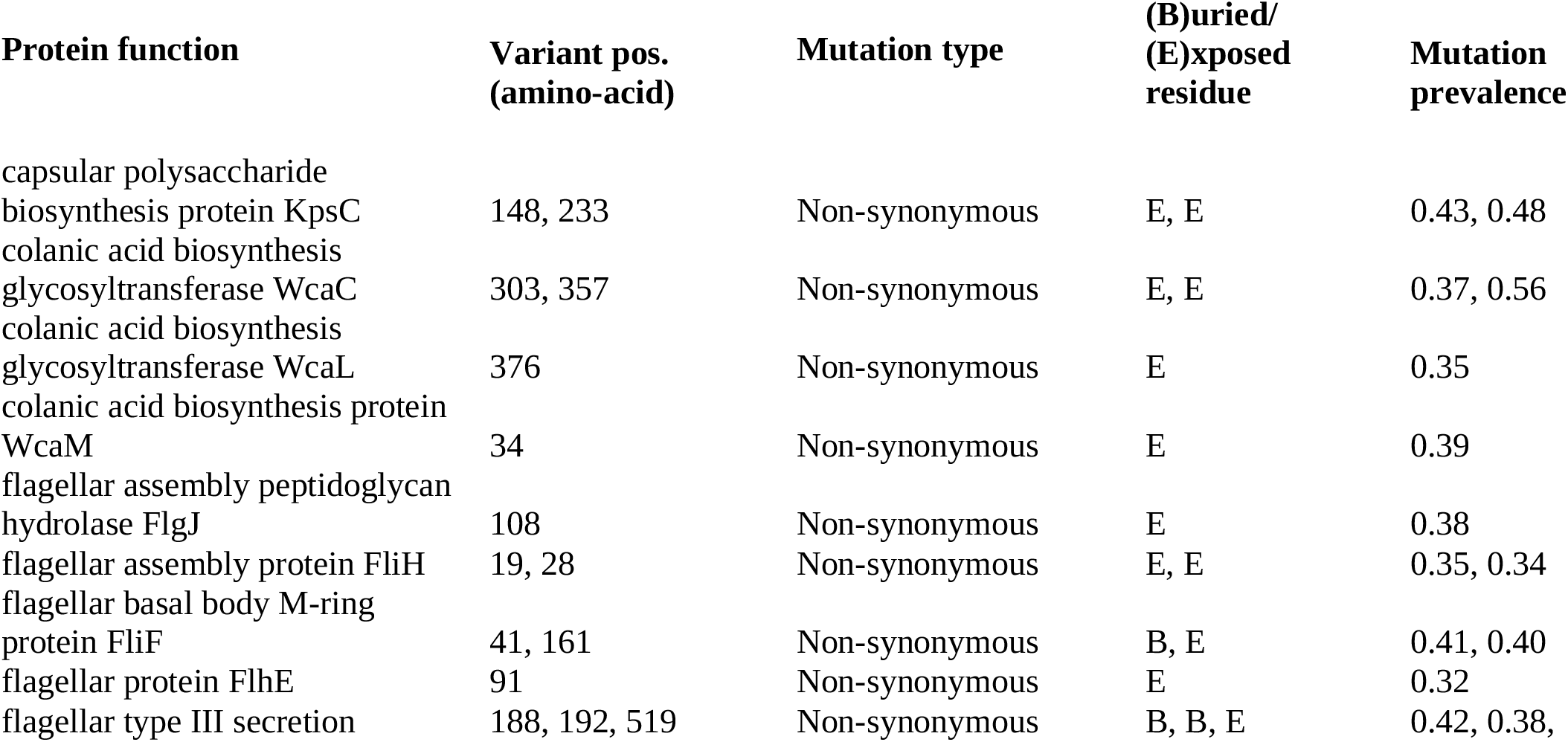
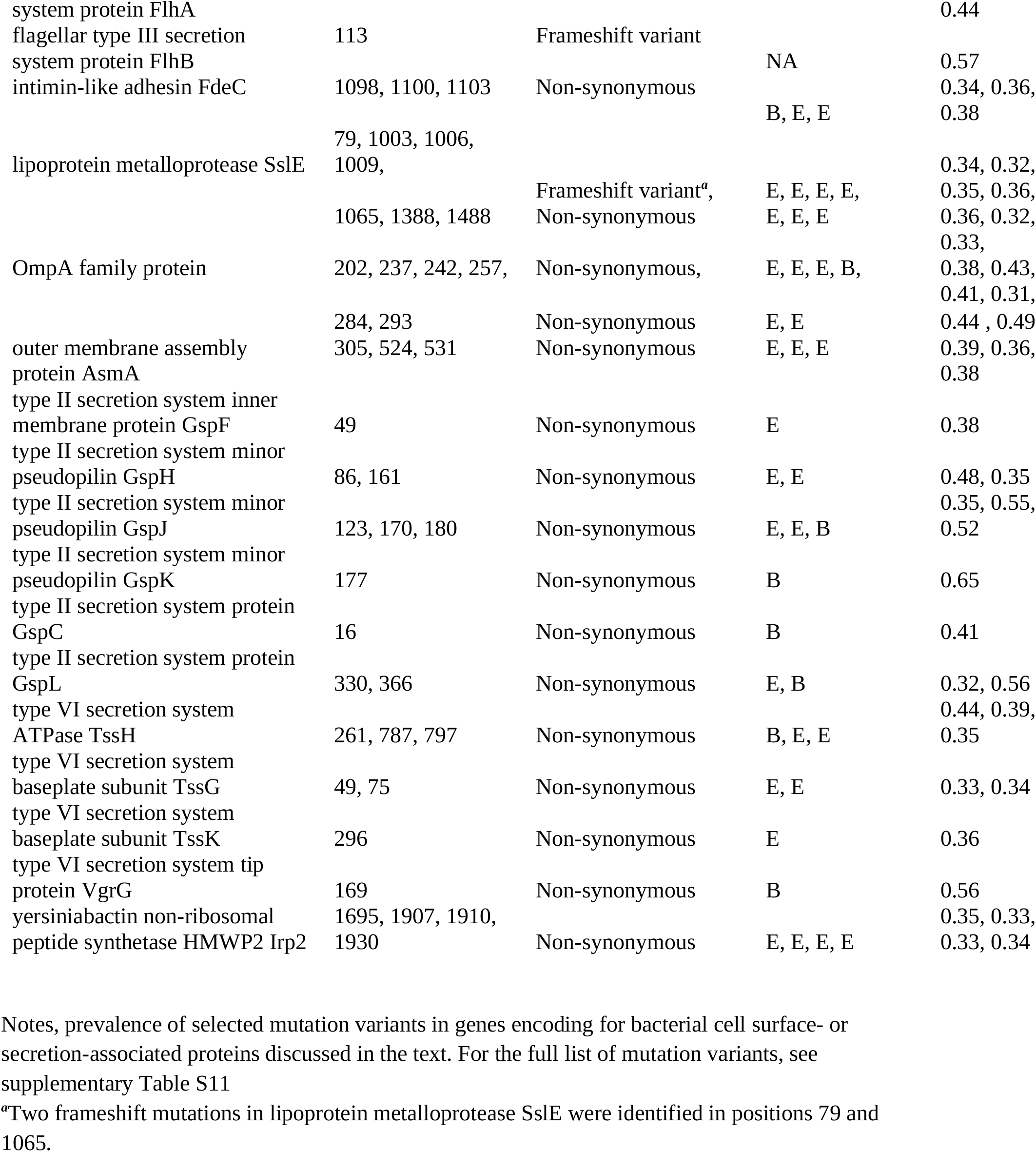
Mutational analysis of functional genes reveals IBD-specific adaptations

Lastly, several of the mutated genes we found in clade-III also appeared with mutation variants in other clades (supplementary Table S12), although the positions of the variants were different. This may suggest a convergent evolution process, and possible adaptations of the *E. coli* strains, regardless of the clade, to the intestines of patients with IBD.

## DISCUSSION

Comparative genomic studies of *E. coli* in IBD have focused mostly on AIEC strains isolated from mucosal samples of patients with CD, and were unable to define genomic markers associated with this phenotype [^19,21^]. Analyses of commensal *E. coli* from fecal and aspirate samples of patients with IBD could not identify IBD-specific strains, genes, or alleles [^22-24^]. Here we characterized large and diverse collections of *E. coli* genomes from different IBD phenotypes to obtain a broader understanding of this species in the context of IBD pathogenesis.

We could classify these diverse *E. coli* strains into five distinct clades. Although no clade was unique to IBD, clade-III (corresponding to the B2 phylogroup) was more prevalent in UC (49%) and healthy controls (41%) than in patients with CD (31.5%) or pouchitis (17.5%). This finding corroborates reports observing a comparable prevalence of *E. coli* isolates of 47% to 55% in biopsy tissues of patients with UC [^58,59^]. We did not detect associations between the relative abundance or the clade affiliation of *E. coli* and intestinal inflammation, in contrast to ^60^Mirsepasi-Lauridsen et al. that found that patients with active UC colonized with *E. coli* B2 had significantly increased levels of fecal calprotectin.

The genomic metabolic capacities varied across the clades and IBD phenotypes. Of particular interest were the sialidase genes, that encode enzymes that cleave sialic acid from sialylated mucins, and were exclusively present in clade-III. Strains from this clade may therefore be better-adapted for utilizing host mucin glycans. Sialylated glycans increase from the ileum to the colon along the gut [^61^]. As patients after pouch surgery are missing a colon and the pouch is made from ileal tissue, this might explain the observation of fewer *E. coli* strains capable of utilizing sialylated mucins in these patients.

In contrast to a study that found a higher abundance of *E. coli pduC* gene in the microbiome of patients with CD compared to healthy subjects [^51^], we observed comparable prevalence of this gene in all IBD phenotypes and in healthy subjects. The *pdu* operon (propanediol) is part of a pathway involved in the metabolism of fucose. It was found that utilization of 1,2-propanediol by AIEC derived from patients with CD is critical for promoting T cell-driven intestinal inflammation in a mouse colitis model [^51^] and is correlated with increased cellular invasion and persistence in vitro [^62^]. The ability to metabolize fucose together with sialic acid (two major components of intestinal mucus) is linked with the expansion of enteropathogenic species like *Salmonella typhimurium* and *Clostridium difficile* which exploit increased mucosal carbohydrate availability following disruption of commensal microbiota by antibiotics [^63^]. On the other hand, the *E. coli* strains found in patients with a pouch were enriched in functions for the utilization of the fructose-containing polysaccharide inulin. Inulin supplementation was shown to reduce endoscopic and histologic inflammation of the mucosa coupled with increased butyrate concentration in patients with a pouch [^64^].

Here we go beyond preliminary reports of higher within-host growth rates of *E. coli* in CD [^42^] into a comprehensive analysis of the major three IBD phenotypes. Enigmatically, patients with a pouch harbor *E. coli* with the highest growth rates, even in the presence of antibiotics. Such high growth rates may correspond to higher levels of nitrate that accumulate in the inflamed gut that allow *E. coli* to perform anaerobic respiration rather than fermentation and consequently, grow faster [^65^]. Nevertheless, we did not observe an association between the iRep and calprotectin levels of the patients.

The Clade-III strains encoded the highest number of putative virulence factors, including *α*-hemolysin, serine-protease autotransporters, the yersiniabactin iron uptake system and the island for biosynthesis of the genotoxic molecule colibactin, confirming previous reports on B2 phylogroup strains [^20,12^]. Recently, mutational signatures that are caused by colibactin exposure were identified in human organoids and matched the mutational signature of human colorectal cancer tumors, implying that colibactin exposure increases colorectal cancer risk [^66^]. Conversely, the *pks* genomic island is also required for the anti-colitis properties of the probiotic *E. coli* strain Nissle 1917 [^67^] that has shown clinical benefit in maintaining remission in patients with UC [^68^]. Using publicly available metagenomic data, we previously showed that the prevalence of *E. coli* colibactin genes is similar between IBD and non-IBD controls, but the relative abundance of colibactin *pks* island is threefold higher in IBD [^69^]. Here we show that *E. coli* from patients with UC are twice more likely to encode the *pks* island than strains from patients with CD or pouch. Combined with the direct effect of years of chronic inflammation of the colon that characterizes UC, these patients may thus be further exposed to a higher risk of developing colorectal cancer, and should undergo surveillance colonoscopy, even during years of remission.

When non-synonymous mutations occur independently in the same gene in multiple strains, this suggests that the gene in question is involved in host adaptation [^70^]. Here, we have identified multiple IBD-specific non-synonymous and frameshift mutations in *E. coli* genes related to bacterial cell envelope and secretion components. This could reflect adaptation to the overly-active host immune response [^2^], that enables such *E. coli* strains to expand in the gut of patients with IBD [^5,8,9^]. Some mutations could actually make IBD-associated strains less proinflammatory to the host. A striking example is the numerous mutations we detected in the metalloprotease SslE (including two frameshift mutations) and the type II secretion system that exports it [^71^]. SslE is generally involved in *E. coli* degradation of mucin substrates and colonization of the intestinal and urinary tract [^72^], in addition to stimulating the production of reactive nitrogen and oxygen species and proinflammatory chemokines in mouse macrophages [^55^]. Thus, loss of SslE function is expected to reduce the pathogenicity of strains that encode it to their hosts. Some IBD-associated mutations observed in envelope proteins such as OmpA, which are known to be highly immunogenic [^53^], could also result in less inflammatory variants of these bacterial proteins. Indeed, we observed that over 80% of the mutations occurred at the surface-exposed regions of those *E. coli* proteins, and can thus affect immunogenicity.

*E. coli* is often considered as an opportunistic pathogen in IBD, but direct mechanistic links between its potential virulence and inflammation *in-vivo* (in patients with IBD) are still missing. Collectively, the current study presents *E. coli* not as a pathogen, but rather as a commensal species that is better adapted to the IBD gut and can benefit from intestinal inflammation. Furthermore, the evidence presented here indicates evolution toward lower virulence in some of the strains, which could explain the lower proinflammatory potential previously observed in *E. coli* strains from patients with pouchitis [^26^]. These results may have important clinical implications, they show that the inflammatory milieu primes the expansion of specific bacteria, such as *E. coli*, rather than these bacteria triggering inflammation and that these bacteria grow rapidly in the presence of antibiotics.

## Acknowledgments

The authors wish to thank the members of the RMC IBD Center and the Comprehensive Pouch Clinic for their help and stimulating discussion of the project. The authors wish to thank Stefan Green of the University of Illinois at Chicago, for his continued expert help in the metagenomic sequencing.

## Ethics approval and consent to participate

The study was approved by the local institutional review board (0298–17) and the National Institutes of Health (NCT01266538). All patients signed informed consent before inclusion to the study.

## Contributors

V.D. U.G. and I.D. conceived and designed the study; V.D. developed the bioinformatic analysis and the comparative genomic pipelines and analyzed the metagenomic data; L.R. analyzed the metagenomic data; K.R. and I.D. collected, analyzed and provided the clinical data; N.W. and I.D. enrolled and examined the patients; V.D. U.G. and I.D. wrote the paper. All authors read, discussed, and approved the final manuscript.

## SUPPLEMENTAL MATERIAL TABLES

**Supplementary Table S1** - The *E. coli* genomes dataset and subjects’ clinical metadata

**Supplementary Table S2** - The *E. coli* MAGs assembly quality controls statistics

**Supplementary Table S3** - Genes with KEGG Orthologs (KO) annotations that significantly differed in prevalence between the *E. coli* clades

**Supplementary Table S4** - Carbohydrate-active enzyme (CAZy) families that significantly differed in prevalence between the *E. coli* clades

**Supplementary Table S5** - Virulence factor genes (based on VFDB) that significantly differed in prevalence between the *E. coli* clades

**Supplementary Table S6** - Antibiotic resistance genes (based on CARD) that significantly differed in prevalence between the *E. coli* clades

**Supplementary Table S7**- KO in *E. coli* strains from clade-III that differed significantly in prevalence between the subject clinical phenotypes

**Supplementary Table S8** - CAZy families in *E. coli* strains from clade-III that differed significantly in prevalence between the subject clinical phenotypes

**Supplementary Table S9** - Virulence genes in *E. coli* strains from clade-III that differed significantly in prevalence between the subject clinical phenotypes

**Supplementary Table S10** - Antibiotic resistance genes (based on CARD) in *E. coli* strains from clade-III that differed significantly in prevalence between the subject clinical phenotypes

**Supplementary Table S11** - Non-synonymous mutations in functionally-annotated genes of *E. coli* strains from clade-III

**Supplementary Table S12** - Mutated genes found in clade-III that appeared with mutation variants in the other clades

## SUPPLEMENTARY MATERIALS AND METHODS

### Pipeline for the reconstruction and analysis of genomes from metagenomes

Metagenome samples where the *E. coli* MAGs failed to pass our minimum accepted quality criteria (completeness ≥75% and contamination **≤**5%), were *de-novo* assembled with metaSPAdes v3.14 [^1^Nurk et al. 2017] using default parameters and ‘--meta’ option, including the read error corrector BayesHammer module, instead of the Unicycler assembler pipeline. The resulting contigs were used for binning using MetaBAT2 v2.15 [^2^Kang et al. 2019] with ‘-m 1500’ option (minimum size of a contig for binning which should be ≥1500) to obtain *E. coli* genome bins. The quality control steps with CheckM were repeated for the newly generated MAGs, and those that passed the accepted quality criteria were added into the analysis pipeline with the rest of the steps unchanged.

### Core genes SNP analysis

Core genes SNP analysis of the *E. coli* genomes was performed using Prokka v1.13.3 [^3^Seemann et al. 2014] in order to obtain the coding genes, and using Roary v3.13 [^4^Page et al. 2015] for pangenome building and a core gene alignment. To convert the resulting core gene alignment fasta file into a SNP distance matrix, the tool SNP-dists (https://github.com/tseemann/snp-dists) was used.

### E. coli phylogroups typing

To perform *in silico* phylogroup typing for the *E. coli* genomes, the ClermonTyping tool (https://github.com/A-BN/ClermonTyping) was used, and the assembled contigs of each genome were used as an input [^5^Beghain et al. 2018].

### Identification of mutation in ciprofloxacin antibiotic target genes

The primary target genes for ciprofloxacin antibiotic, *gyrA* and *parC*, were extracted from the coding genes predicted by Prokka (see above). Point mutations in the genes in positions known to confer resistance were identified as described in ^6^Dubinsky et al. 2020.

